# Aortic regurgitation provokes phenotypic modulation of smooth muscle cells in the normal ascending aorta

**DOI:** 10.1101/2023.02.08.527682

**Authors:** Brittany Balint, Inés García Lascurain Bernstorff, Tanja Schwab, Hans-Joachim Schäfers

## Abstract

**Background:** Aortic complications are more likely to occur in patients with ascending aortic aneurysms and concomitant aortic regurgitation (AR). AR may have a negative impact on the aortic wall structure even in patients with tricuspid aortic valves and absence of aortic dilatation. It is unknown whether smooth muscle cell (SMC) changes are a feature of AR-associated aortic remodeling.

**Methods:** Non-dilated aortic samples were harvested intra-operatively from individuals with normal aortic valves (n=10) or those with either predominant aortic stenosis (AS; n=20) or AR (n=35). Tissue from each patient was processed for immunohistochemistry or used for the extraction of medial SMCs. Tissue and cells were stained for markers of SMC contraction (alpha-smooth muscle actin; ASMA), synthesis (vimentin) and senescence (p16/p21). Replicative capacity was analyzed in cultured SMCs from AS- and AR-associated aortas. A sub-analysis compared SMCs from individuals with either TAVs or BAVs to rule out the effect of aortic valve morphology.

**Results:** In aortic tissue samples, AR was associated with decreased ASMA and increased vimentin, p16 and p21 compared to normal aortic valves and AS. In cell culture, SMCs from AR-aortas had decreased ASMA and increased vimentin compared to SMCs from AS-aortas. AR-associated SMCs had increased p16 and p21 expression, and they reached senescence earlier than SMCs from AS-aortas. In AR, SMC changes were more pronounced with the presence of a BAV.

**Conclusions:** AR itself negatively impacts SMC phenotype in the ascending aortic wall, which is independent of aortic diameter and aortic valve morphology. These findings provide insight into the mechanisms of AR-related aortic remodeling, and they provide a model for studying SMC-specific therapies in culture.

## Introduction

Aneurysms of the ascending aorta are frequently associated with life-threatening complications, i.e., aortic dissection, rupture, and sudden death. These aneurysms may derive from multiple origins, including connective tissue disorders with monogenetic causes (i.e., Marfan and Loeys-Dietz syndromes)^1, 2^ or bicuspid aortic valves (BAV)^3^, in which the etiology is less defined. Both in aneurysms due to connective tissue or BAV disease, aortic valve regurgitation (AR) is a risk factor for aortic complications^4^.

There are common features of degenerative aortic wall remodeling in ascending aortic aneurysms, including extracellular matrix (ECM) degeneration^3, 5, 6^. The underlying mechanisms are incompletely understood. BAVs are associated with aortic stenosis (AS) and AR, and turbulent blood flow across the BAV is a significant cause of aneurysm formation^7–9^. Irrespective of aortic valve morphology, AS is associated with turbulent blood flow, whereas AR is associated with increased stroke volume, both resulting in subsequent increases in shear stress^10, 11^. Interestingly, the BAV-associated ascending aorta dilates faster when AR is present^12^, and the probability of aortic complications is higher with AR compared to normally functioning valves and AS^13^. Less data are available for ascending aortic aneurysms in the absence of a BAV. We recently studied non-dilated ascending aortas in individuals with tricuspid aortic valves (TAVs). We found that AR was associated with a higher degree of aortic wall remodeling compared to AS or normally functioning valves, which included elastin degradation and mucoid ECM accumulation in the medial layer^14^.

Smooth muscle cells (SMCs) are an important component of the aortic media. Contractile SMCs are embedded in the ECM, and they maintain structural and functional integrity of the aorta by circumferentially wrapping around the vessel to impart coordinated vascular tone. Previous histologic studies have identified phenotypic modulation of SMCs in aneurysmal aortas, whereby SMCs lose their contractile properties and switch to a synthetic phenotype^15^. SMCs with synthetic properties are crucial during development and in response to injury, but they negatively impact the adult aortic wall by secreting ECM degrading enzymes^16^. Others have identified non-contractile senescent SMCs in the aneurysmal ascending aorta, which are capable of degrading surrounding ECM through their seno-destructive secretory phenotype^6^. The mechanisms by which SMCs switch from the contractile phenotype to maladaptive synthetic or senescent states in the vulnerable ascending aortic wall remains uncertain. Even less is known regarding the role of medial SMCs in aortic wall remodeling in relation to AR.

We hypothesized that different flow characteristics due to aortic valve dysfunction could alter SMC phenotypes in the ascending aorta. We first histologically assessed medial cell phenotypes in the aorta of individuals with and without aortic valve dysfunction. We then isolated ascending aortic SMCs from individuals with either predominant AS or AR, and studied their phenotypic properties in culture. We exclusively studied SMCs from non-dilated ascending aortas to isolate our findings from potential confounding causes of aortic wall remodeling. To rule out the effect of aortic valve morphologies, we compared SMCs from individuals with either TAVs or BAVs.

## Methods

The data presented in this manuscript are available upon reasonable request from the corresponding author. This study complies with the Declaration of Helsinki, and was carried out with approval from the regional ethics committee (Ständige Ethikkommission der Ärztekammer des Saarlandes, Proposal # 47/14). With the exception of donor controls, written informed consent was obtained from all patients. For donor samples, potential identifying data and characteristics were excluded.

### Patient Enrollment and Exclusion Criteria

Fifty-five consecutive cardiac patients undergoing aortic valve surgery for either predominant aortic stenosis (AS; n=20) or aortic regurgitation (AR; n=35) were enrolled in this study. For controls, aortic tissue was collected from autopsy samples (n=10). AR patients were included only if the AR was due to cusp causes (i.e., prolapse, retraction), and if their aortic root dimensions were within the normal range. Aortic valve morphology was determined pre-operatively by either trans-thoracic or trans-esophageal echocardiology, and was confirmed intra-operatively by the primary surgeon. For this study, only patients with tricuspid aortic valves (TAV) were included. Prior to surgery, aortic dimensions were determined by computed tomography, and they were confirmed intra-operatively via trans-esophageal echocardiography. Aortic diameters measuring ≥40mm were considered dilated^17^, and were therefore excluded from this study. Patients were excluded if they had any serological evidence of chronic viral diseases (i.e., HIV, Hepatitis B or C) or if they showed any clinical signs of connective tissue disorders (i.e., Marfan or Loeys-Dietz syndrome). Aortic samples were macroscopically analyzed and those with evidence of inflammatory disease (i.e., atherosclerosis, aortitis) were also excluded. For the control group, aortic tissue samples were collected from autopsies of individuals without any macroscopic evidence of cardiac valve disease or aortic dilatation.

### Procurement of ascending aortic tissue

For each of the AS and AR patients, a circumferential fragment of aortic tissue (4-5mm width) was excised from the anterior circumference of the thoracic aorta, 5-10mm above the sinotubular junction. The excised tissue was divided in the operating room, and a portion was immediately fixed in 4% phosphate-buffered formalin for histological studies. The remaining aortic tissue was carefully transferred in PBS to a sterile tissue culture hood for immediate extraction of medial SMCs. For autopsy samples, full circumferential aortic rings were excised and immediately fixed in 4% phosphate-buffered formalin. Due to the nature of tissue retrieval from autopsy samples, fresh tissue was unavailable for SMC isolation.

### SMC Isolation and culture

SMCs were isolated from the medial layer of aortic tissue samples through enzymatic digestion with Liberase™, a purified collagenase blend. First, to ensure that only medial cells would be isolated, we separated the medial layer from the aortic tissue. This was done by gently scraping away the endothelial cell layer with a scalpel, and then peeling away the adventitia and the outermost medial layers with forceps. The remaining medial tissue fragment was washed in PBS and then cut into ~1×1mm pieces to be used for the enzymatic digestion.

Medial tissue fragments were added to 1.5mL Eppendorf tubes containing the enzymatic digestion media (830μl of M199 Media (ThermoFisher Scientific, 11150059; + 0.5% FBS + 1% penicillin/streptomycin), 150μl of Liberase™ (Roche, 05401020001) and 30μl of DNAse). The sample was incubated for 2h (37°C, 5% CO_2_, 95% humidity), and then the digested supernatant was collected through a cell strainer and maintained on ice until the end of the final digestion. The undigested aortic medial tissue was transferred to a new 1.5mL Eppendorf tube containing fresh enzymatic digestion media. The digestion process was repeated for a second round with an incubation period of 1.5h. The collected supernatant from both digestions were combined and then centrifuged at 750 RPM for 6 minutes at 4°C. The resulting cell pellet was washed with PBS, and then reconstituted in M199 media containing 10% FBS and 1% penicillin/streptomycin. The cells were plated on 0.4% gelatin-coated 60 mm culture dishes (Gelatin Type B Powder, Sigma-Aldrich, G9391; ThermoFisher Scientific Cell Culture Petri Dishes, 150340), and incubated at 37°C, 5% CO_2_, and 95% humidity. Media was changed 24h after the SMC isolation was completed, and then every 48h until confluence was reached. At this point, the cells were detached from the culture dishes (Trypsin/EDTA solution, ThermoFisher Scientific, R001100) and were either re-plated on coverslips for immunocytochemistry (cell passage 1), or were re-plated on fresh gelatin-coated dishes, repeatedly until senescence was reached. The maximum cell passage achieved by the SMC culture from each patient was deemed as the replicative capacity. For immunocytochemistry, the SMCs were serum-starved (0.5% FBS) once 80% confluence was reached. The cells were fixed 72h post serum-withdrawal with 4% paraformaldehyde.

### Immunostaining of aortic tissue and cultured SMCs

Formalin-fixed aortic tissue samples were embedded in paraffin, and were sectioned at 1μm thickness. Aortic tissue sections and paraformaldehyde-fixed aortic SMCs were immunolabeled with rabbit polyclonal antibodies against either alpha-smooth muscle actin (ASMA; 1:100, ab5694, Abcam), a contractile protein, or vimentin (1:100, ab137321, Abcam), an intermediate filament protein that is abundantly expressed in synthetic SMCs^18^. Cellular senescence was evaluated in aortic tissue and SMCs by immunolabelling for cyclin-dependent kinase inhibitors, p16^INK4a^ (monoclonal mouse, 1:50, MA5-17054, Invitrogen) and p21^Cip1^ (1:50, MA1-33926, Invitrogen), which demarcate two core senescence pathways^19, 20^. Bound primary antibodies were visualized with fluorescently conjugated secondary goat antibodies targeted against rabbit or mouse (Alexa-594). Aortic tissue samples were counter-stained with DAPI and then were mounted onto microscopy slides. SMC-containing coverslips were mounted onto microscopy slides with a DAPI-containing mounting media (Vectashield®, H-1200-10).

### Microscopy and Image Analysis

Fluorescently labeled aortic tissue samples and SMCs were imaged with a laser scanning confocal microscope (Zeiss LSM, Plan Apochromat). Aortic tissue samples were imaged at 40x with a 1.3 oil objective, and SMCs were imaged at 20x with a 0.8 M27 objective. For each patient, 5-10 regions of interest were captured per stain for both aortic tissue and SMC samples. The fluorescence intensity of ASMA and vimentin were measured in ImageJ (NIH) and normalized to the background intensity of each image. The percent positivity of p16 and p21 were also measured in ImageJ by counting the number of positive nuclei in each region of interest, and normalizing by the total number of nuclei in the image.

### Statistics

All statistical analyses were carried out using Prism 9 (Graphpad Software). All datasets were tested for normality using the D’Agostino and Pearson omnibus test. In the case of normally distributed data, comparisons between groups were made with the Student’s t-test or the one-way ANOVA with the Bonferroni-adjusted post-hoc test. If at least one data set did not pass the normality test, data sets were compared with the Mann-Whitney U test or the Kruskal-Wallis test with Dunn’s multiple comparisons post-hoc test. Linear regression analyses were used to assess for relationships between continuous variables. Patient age data are presented as mean ± standard deviation. Cell passage data are presented as median ± standard deviation. Statistical significance was set at *P*<0.05.

## Results

### Patient Characteristics

Ascending aortic tissue was collected intra-operatively from 55 consecutive patients undergoing aortic valve surgery for either predominant AS (n=20) or predominant AR (n=35). Control ascending aortic tissue was collected from autopsy samples with no evidence of aortic valve disease or dilatation (n=10). The mean age of patients with predominant AS was 57.9±11.7 (min.:19, max.:70) years, and in patients with predominant AR it was 59.4±13.8 (min.:15, max.:84) years. Patients from the control group were significantly younger than those from the AS and the AR groups (mean: 39.7±6.1, min.: 28, max.: 50 years; *P*=0.0001). There was no significant difference in age between the AS and AR groups (*P*=0.89). Aortic dimensions were similar between AS and AR groups (aortic sinus: *P*=0.10, sinotubular junction: *P*=0.23, mid-ascending aorta: *P*=0.33). Due to the nature of tissue retrieval from autopsy samples, in vivo patient data were not available for the control group. Clinical characteristics are presented in **Table 1**.

**Table 1.** Patient Characteristics.

|  | Control | Aortic Stenosis | Aortic Regurgitation |
| --- | --- | --- | --- |
| # of Patients | 10 | 20 | 35 |
| Patient Age (Range) | 28 – 50 | 19 – 70 | 15 – 84 |
| Patient Age (Mean±SD) | 39.7±6.1* | 57.9±11.7 | 59.4±13.8 |
| Ascending Aortic Diameter (Range; cm) | — | 2.4 – 3.9 | 2.6 – 4.0 |
| Ascending Aortic Diameter (Mean±SD) | — | 3.2±0.5 | 3.4±0.4 |
|  | Co-Morbidities |  |  |
| Hypertension (%) | — | 6 (30) | 10 (29) |
| Hyperlipidemia (%) | — | 5 (25) | 11 (31) |
| Smoker (%) | — | 3 (15) | 5 (14) |
| Diabetes (%) | — | 1 (5) | 2 (6) |
|  | Medications |  |  |
| β-Blocker (%) | — | 4 (20) | 4 (11) |
| ACE Inhibitor (%) | — | 2 (10) | 7 (20) |
| Diuretic (%) | — | 3 (15) | 4 (11) |
| Calcium Channel Blocker (%) | — | 1 (5) | 5 (14) |
| Statin (%) | — | 2 (10) | 3 (9) |

SD=standard deviation, ACE=angiotensin converting enzyme, \* $P<0.05$

For a sub-analysis, AS and AR patients were further divided into groups based on aortic valve morphology. Of the patients with predominant AS, 11 had a TAV and 9 had a BAV. There was no significant difference in patient age between TAV and BAV subgroups (*P*=0.07). In the AR group, 10 patients had a TAV and 25 had a BAV. The AR patients with a BAV were significantly younger than those with a TAV (*P*=0.02). Clinical characteristics based on aortic valve morphology are presented in **Table 2.**

**Table 2.**
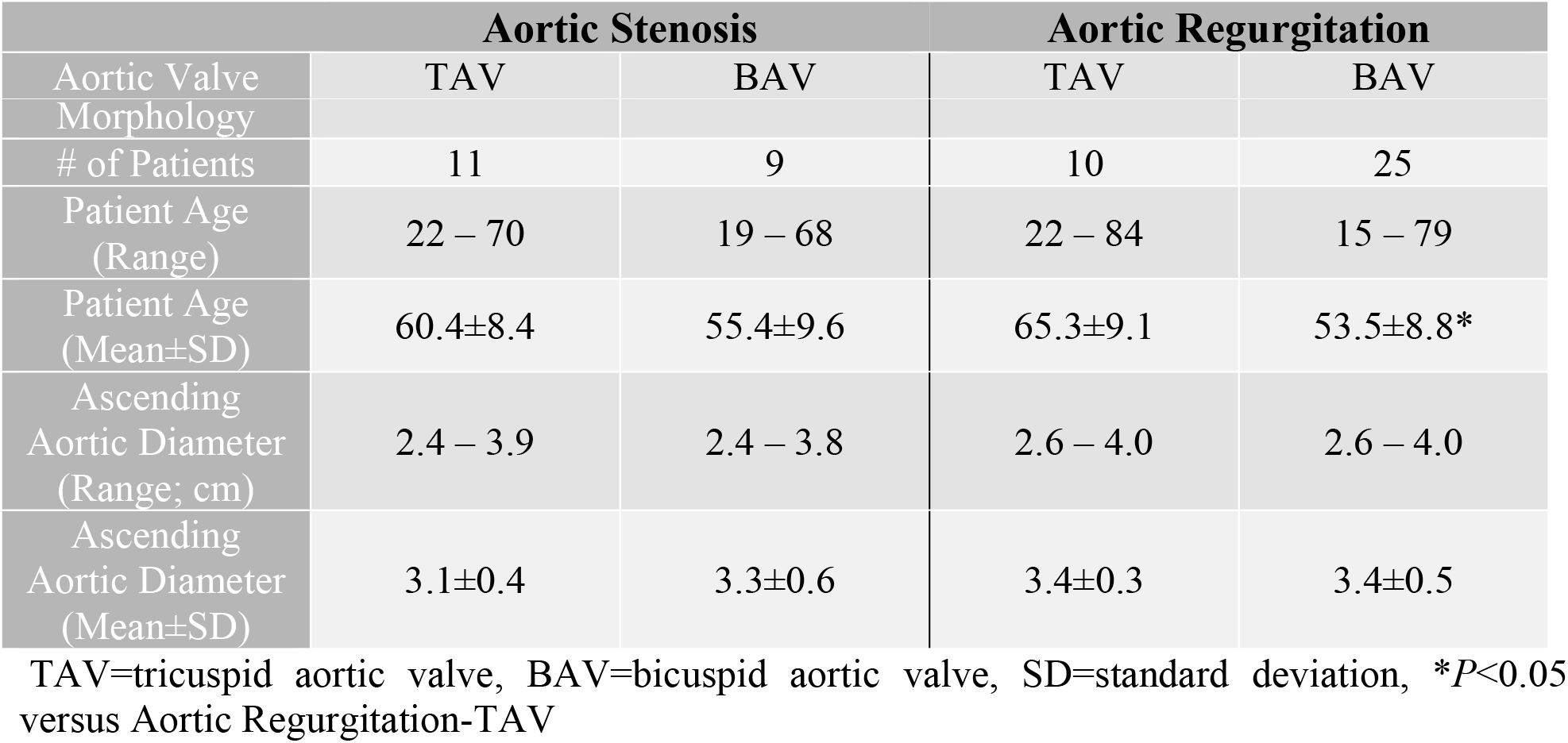
Clinical Characteristics Based on Aortic Valve Morphology.

### AR is associated with actin loss and increased vimentin in the ascending aortic wall

We first aimed to determine whether phenotypic switching of SMCs is a feature of aortic valve disease-associated aortic remodeling. Non-dilated ascending aortic tissue was assessed for the expression of ASMA (contractile phenotype) and vimentin (synthetic phenotype). The expression of ASMA was decreased in AR-associated aortas compared to AS-associated aortas (*P*=0.03) and control aortas (*P*<0.0001; **Figure 1a,b**). Conversely, the expression of vimentin was increased in AR-associated aortas compared to AS-associated aortas and control aortas (all *P*<0.0001; **Figure 1a,b**). ASMA was decreased in AS-associated aortas compared to control (*P*=0.03), and there was a trend towards increased vimentin (*P*=0.06; **Figure 1a,b**). Assessing for a switch from contractile to synthetic SMCs, linear regression analyses revealed that ASMA expression negatively correlated with vimentin expression within aortic samples for all three groups (Control: R^2^=0.42, *P*=0.04, AS: R^2^=0.29, *P*=0.01, AR: R^2^=0.4, *P*<0.0001; **Figure 1c**). Furthermore, ASMA expression inversely correlated with patient age for all three groups (Control: R^2^=0.45, *P*=0.03, AS: R^2^=0.35, *P*=0.007, AR: R^2^ =0.4, *P*<0.0001; **Figure 1d**). Vimentin expression positively correlated with patient age for the AR group (R^2^=0.36, *P=* 0.0001), while no significant relationship was detected for the control or AS group (Control: R^2^=0.22, *P*=0.69, AS: R^2^=0.13, *P*=0.12; **Figure 1d**).

**Figure 1.**
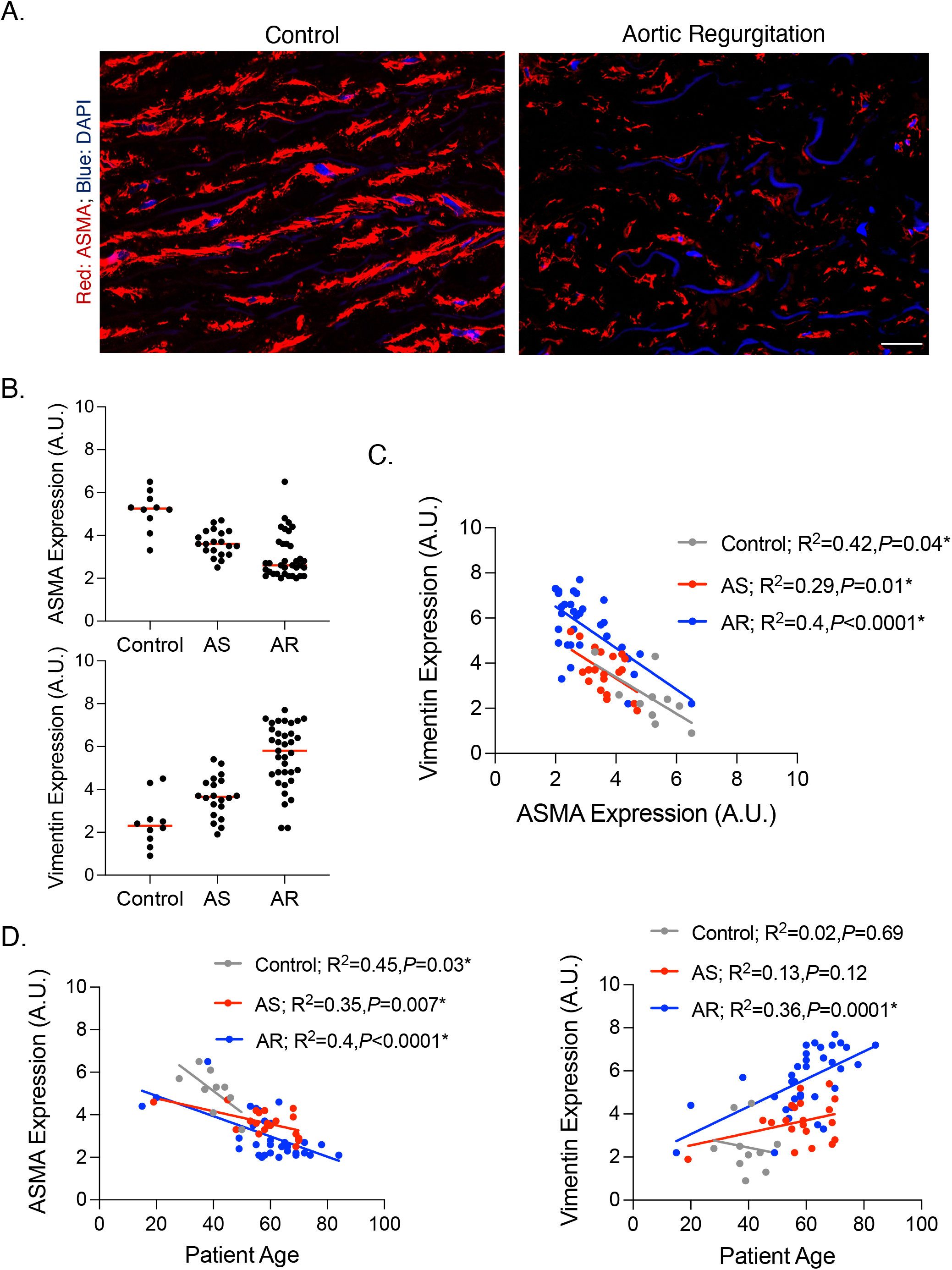
Alpha-smooth muscle actin (ASMA) is decreased while vimentin is increased in aortic regurgitation (AR)-associated aortas. **A.** Fluorescent micrographs of ASMA in control (left) and AR- (right) associated ascending aortic tissue. **B.** Graphs depicting ASMA (top) and vimentin (bottom) expression in the ascending aorta from each group. Horizontal bars represent median values. **C.** Graph depicting the relationship of ASMA and vimentin for each group. **D.** Graphs depicting the relationship between ASMA (left) or vimentin (right) and patient age for each group. A.U.= arbitrary units; *=statistical significance; scale bar = 20 μm.

### AR is associated with cellular senescence in the ascending aortic medial layer

To determine whether cellular senescence is a feature of AS- or AR-associated aortic remodeling, we analyzed ascending aortic tissue for the expression of the cell cycle inhibitors, p16^INK4a^ and p21^Cip1^. The expression of both p16^INK4a^ and p21^Cip1^ was increased in AR-associated aortas compared to AS-associated aortas (*P*<0.0001 and *P*=0.0001, respectively) and controls (*P*=0.0005 and *P*=0.0011, respectively; **Figure 2a,b**). There was no significant difference in either p16^INK4a^ or p21^Cip1^ expression between AS-associated aortas and control aortas (all *P*=0.99; **Figure 2a,b**). There was no significant relationship between p16^INK4a^ and patient age in either group (Control: R^2^<0.0001, *P*=0.95, AS: R^2^=0.17, *P*=0.07, AR: R^2^=0.008, *P*=0.60; **Figure 2c**). On the other hand, p21^Cip1^ expression increased with patient age for both the AS (R^2^=0.3, *P*=0.03) and AR (R^2^=0.16, *P*=0.02) groups (**Figure 2d)**. There was no significant association between patient age and p21^Cip1^ expression for the control group (R^2^=0.009, *P*=0.79; **Figure 2d**).

**Figure 2.**
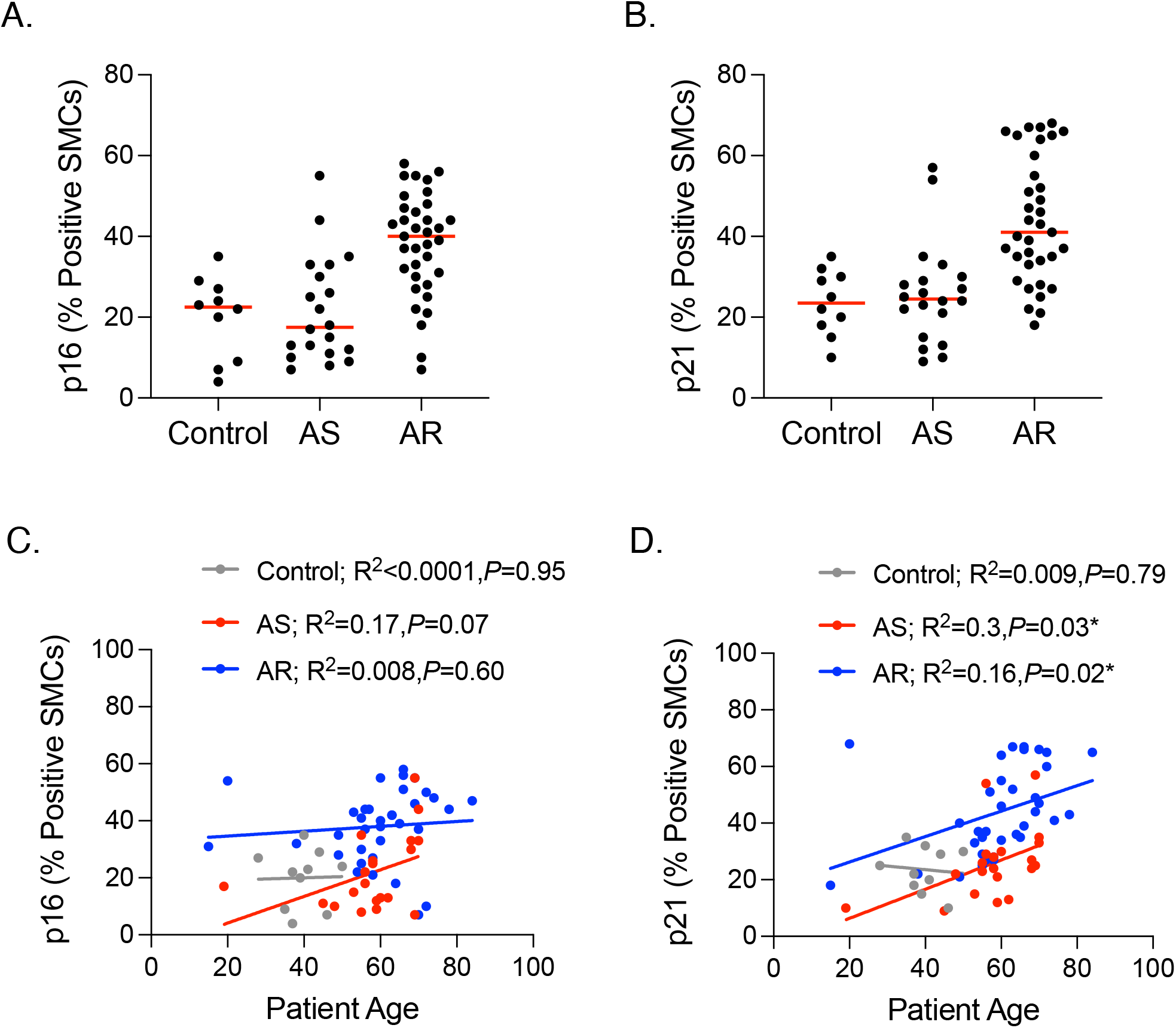
Senescence markers p16^INK4a^ and p21^Cip1^ are increased in aortic regurgitation (AR)-associated aortas. **A-B.** Graphs depicting p16^INK4a^ (**A**) and p21^Cip1^ (**B**) expression in the ascending aorta from control patients, or from patients with aortic stenosis (AS) or AR. Horizontal bars represent median values. **C-D.** Graphs depicting the relationship between p16^INK4a^ (**C**) or p21^Cip1^ (**D**) expression and patient age for each group. SMCs= Smooth muscle cells. *=statistical significance.

### AR-associated changes in the aortic wall are more pronounced in the presence of a BAV

Since BAVs are associated with ascending aortic aneurysms, we next sought to determine whether the presence of a BAV aggravates AS- or AR-associated changes in the aortic wall. Among patients with predominant AS, ASMA expression was decreased in the BAV-associated ascending aorta (BAV-AS) compared to TAV aortas (TAV-AS; *P*=0.003; **Figure 3a**). No significant changes were observed in either the expression of vimentin (*P*=0.96; **Figure 3b**) or the senescence markers when comparing TAV-AS to BAV-AS aortas (p16^INK4a^: *P*=0.17; p21^Cip1^: *P*=0.30; **Figure 3c,d**).

**Figure 3.**
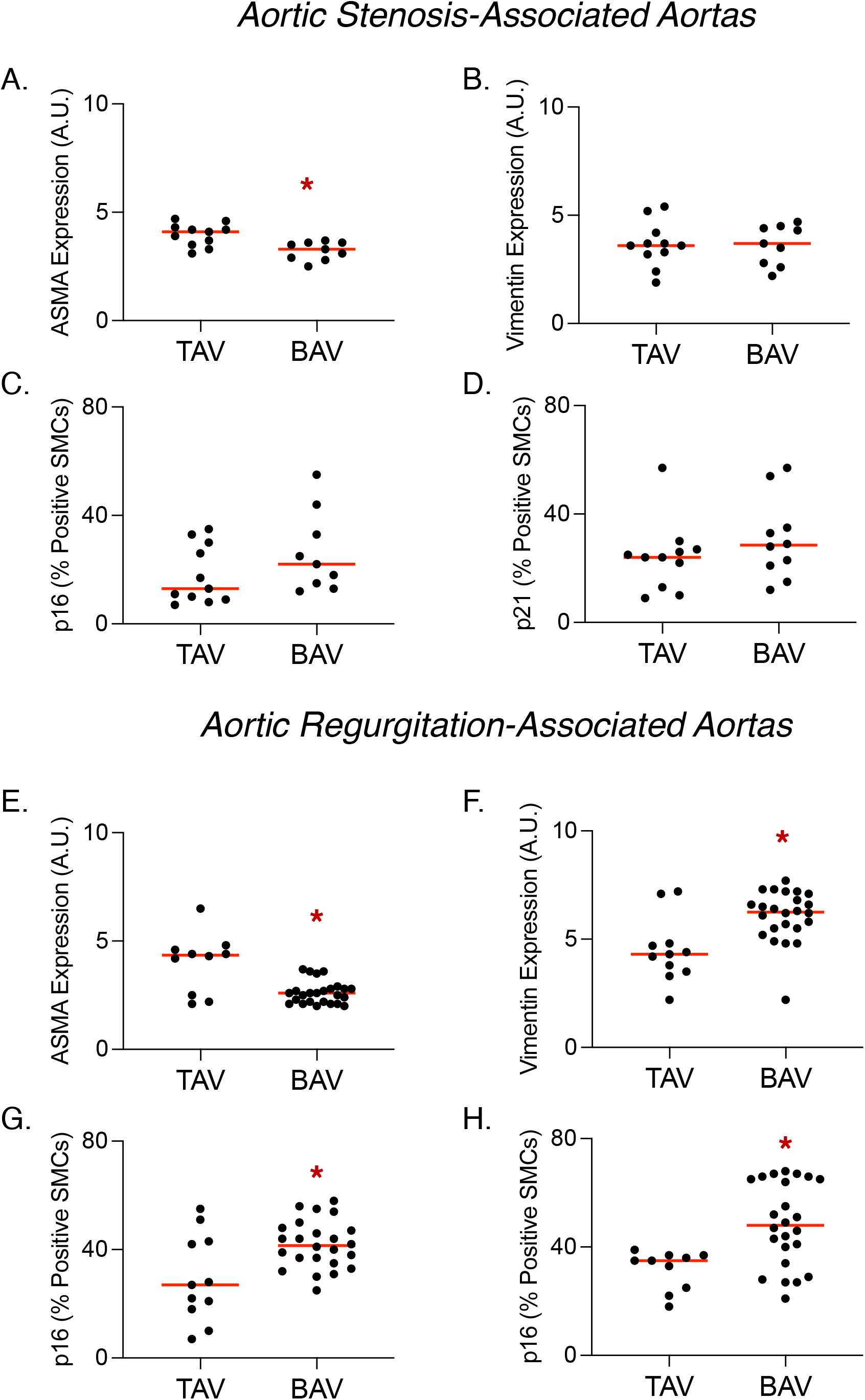
Bicuspid aortic valve (BAV) morphology aggravates phenotypic changes in aortic regurgitation (AR)-associated aortas. **A-D.** Graphs depicting the expression of alpha smooth muscle actin (ASMA; **A**), vimentin (**B**), p16 (**C**) and p21 (**D**) in the aorta of individuals with aortic stenosis and either tricuspid aortic valves (TAV) or BAVs. **E-H.** Graphs depicting the expression of ASMA (**E**), vimentin (**F**), p16 (**G**) and p21 (**H**) in the aorta of individuals with aortic regurgitation and either TAVs or BAVs. SMCs= Smooth muscle cells. Horizontal bars represent median values. *=statistical significance.

For patients with predominant AR, the expression of ASMA was decreased in the ascending aorta of BAV patients (BAV-AR) compared to those with a TAV (TAV-AR; *P*=0.0001; **Figure 3e**). On the other hand, vimentin expression was increased in BAV-AR ascending aortas compared to that of TAV-AR (*P*=0.002; **Figure 3f**). Despite being biologically younger in age, cellular senescence was exacerbated by the presence of BAV, as p16^INK4a^ (*P*=0.005) and p21^Cip1^ (*P*=0.003) expression was increased in BAV-AR aortas compared to TAV-AR aortas (**Figure 3g,h**).

### AR is associated with phenotypic switching of SMCs isolated from the normal ascending aorta

To determine if disruptive SMC changes underlie AR-associated ascending aortic remodeling, we isolated medial SMCs from fresh ascending aortic tissue and studied their phenotypic properties in culture. There was a significant decrease in ASMA expression in SMCs from AR-associated aortas compared to those from AS-associated aortas (*P*=0.0001; **Figure 4a,b**). On the other hand, vimentin expression was increased in AR-associated SMCs versus AS-associated SMCs (*P*<0.0001; **Figure 4a,b**). Similar to our findings in aortic tissue, there was an inverse correlation between ASMA and vimentin expression for both groups (AS: R^2^=0.40, *P*=0.003, AR: R^2^=0.12, *P*=0.02; **Figure 4c**). While patient age increased, ASMA expression decreased (AS: R^2^=0.33, *P*=0.009, AR: R^2^=0.24, *P*=0.003) and vimentin expression increased in SMCs from both groups (AS: R^2^=0.25, *P*=0.02, AR: R^2^=0.24, *P*=0.003; **Figure 4d**).

**Figure 4.**
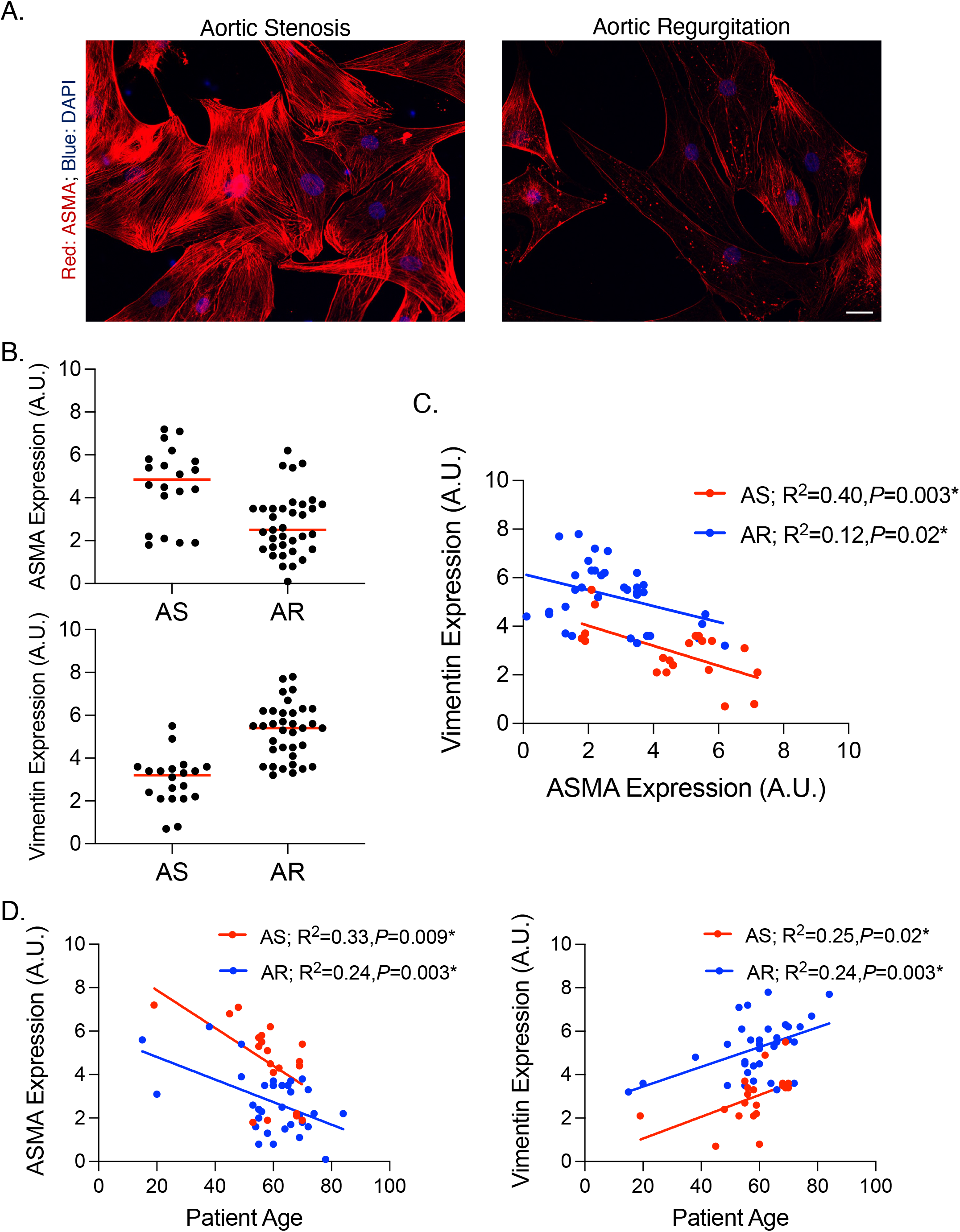
Alpha-smooth muscle actin (ASMA) is decreased while vimentin is increased in smooth muscle cells (SMCs) isolated from aortic regurgitation (AR)-associated aortas. **A.** Fluorescent micrographs of ASMA in aortic stenosis (AS)- (left) and AR- (right) associated SMCs. **B.** Graphs depicting ASMA (top) and vimentin (bottom) expression in SMCs from from each group. Horizontal bars represent median values. **C.** Graph depicting the relationship between ASMA and vimentin in SMCs for each group. **D.** Graphs depicting the relationship between ASMA (left) or vimentin (right) and patient age in SMCs for each group. A.U.= arbitrary units; *=statistical significance; Scale bar = 20 μm.

### Cellular senescence is accelerated in SMCs isolated from AR-associated ascending aortas

We measured cellular senescence in isolated SMCs by analyzing the expression of p16^INK4a^ and p21^Cip1^, and by assessing the replicative capacity of SMCs in culture. There was a trend towards increased expression of p16^INK4a^ in AR SMCs versus AS SMCs (*P*=0.06; **Figure 5a**). The expression of p21^Cip1^ was significantly increased in SMCs isolated from AR-associated aortas compared to that of AS-associated aortas (*P*=0.006; **Figure 5a**). Patient age correlated with p21^Cip1^ expression in both groups (AS: R^2^=0.25, *P*=0.03, AR: R^2^=0.40, *P*<0.0001), whereas no relation was detected between age and p16^INK4a^ (AS: R^2^=0.09, *P*=0.19, AR: R^2^=0.05, *P*=0.20; **Figure 5b**). By allowing SMC cultures from each patient to grow indefinitely, we found that the replicative capacity of SMCs from AR-associated aortas (cell passage 3.0±1.1) was decreased compared to that of SMCs from AS-associated aortas (cell passage 5.0±1.6; *P*<0.0001; **Figure 5c**). This suggests that AR SMCs enter cellular senescence at an accelerated rate compared to SMCs from AS-associated aortas.

**Figure 5.**
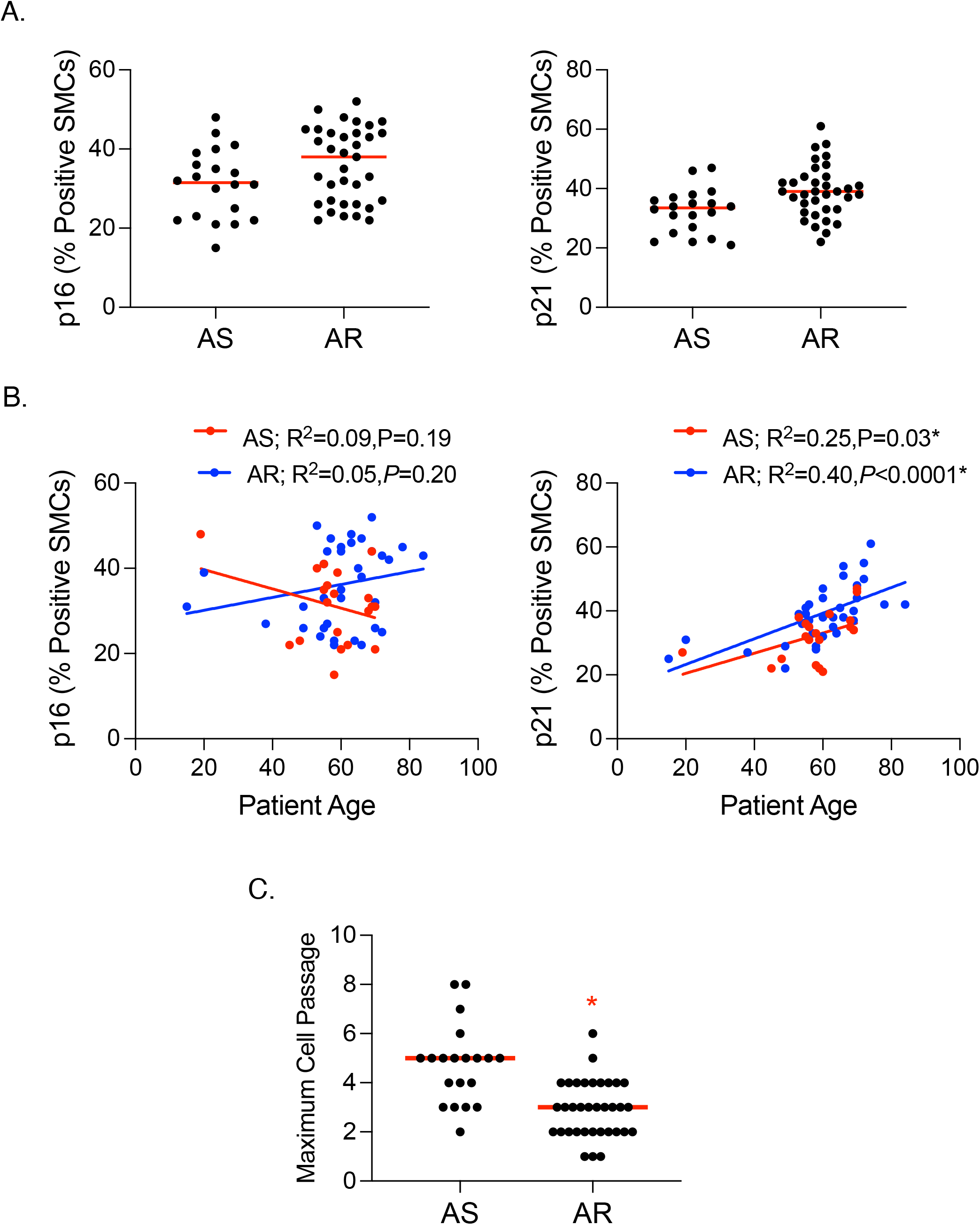
Senescence markers p16^INK4a^ and p21^Cip1^ are increased in smooth muscle cells (SMCs) isolated from aortic regurgitation (AR)-associated aortas. **A.** Graphs depicting p16^INK4a^ (left) and p21^Cip1^ (right) expression in aortic SMCs isolated from patients with aortic stenosis (AS) or AR. **B.** Graphs depicting the relationship between p16^INK4a^ (left) or p21^Cip1^ (right) expression and patient age in SMCs for each group. **C.** Graph depicting the maximum cell passage of cultured SMCs isolated from AS- or AR-associated aortas. Horizontal bars represent median values. *=statistical significance.

### AR-associated phenotypic switching of SMCs is more pronounced in the presence of a BAV

Like in tissue, we assessed phenotypic switching of SMCs from AS- and AR-associated aortas in patients with either a TAV or a BAV. For patients presenting with predominant AS, ASMA expression was decreased in BAV-AS SMCs compared to TAV-AS SMCs (*P*=0.001; **Figure 6a**). The expression of vimentin, however, was increased in BAV-AS SMCs compared to TAV-AS SMCs (*P*=0.02; **Figure 6a**). No differences in SMC senescence were observed between TAV-AS and BAV-AS SMCs, as the expression of p16^INK4a^ and p21^Cip1^ was similar between groups (*P*=0.78 and *P*=0.66, respectively; **Figure 6a**). Furthermore, SMC replicative capacity did not differ between groups (*P*=0.12; **Figure 6b**).

**Figure 6.**
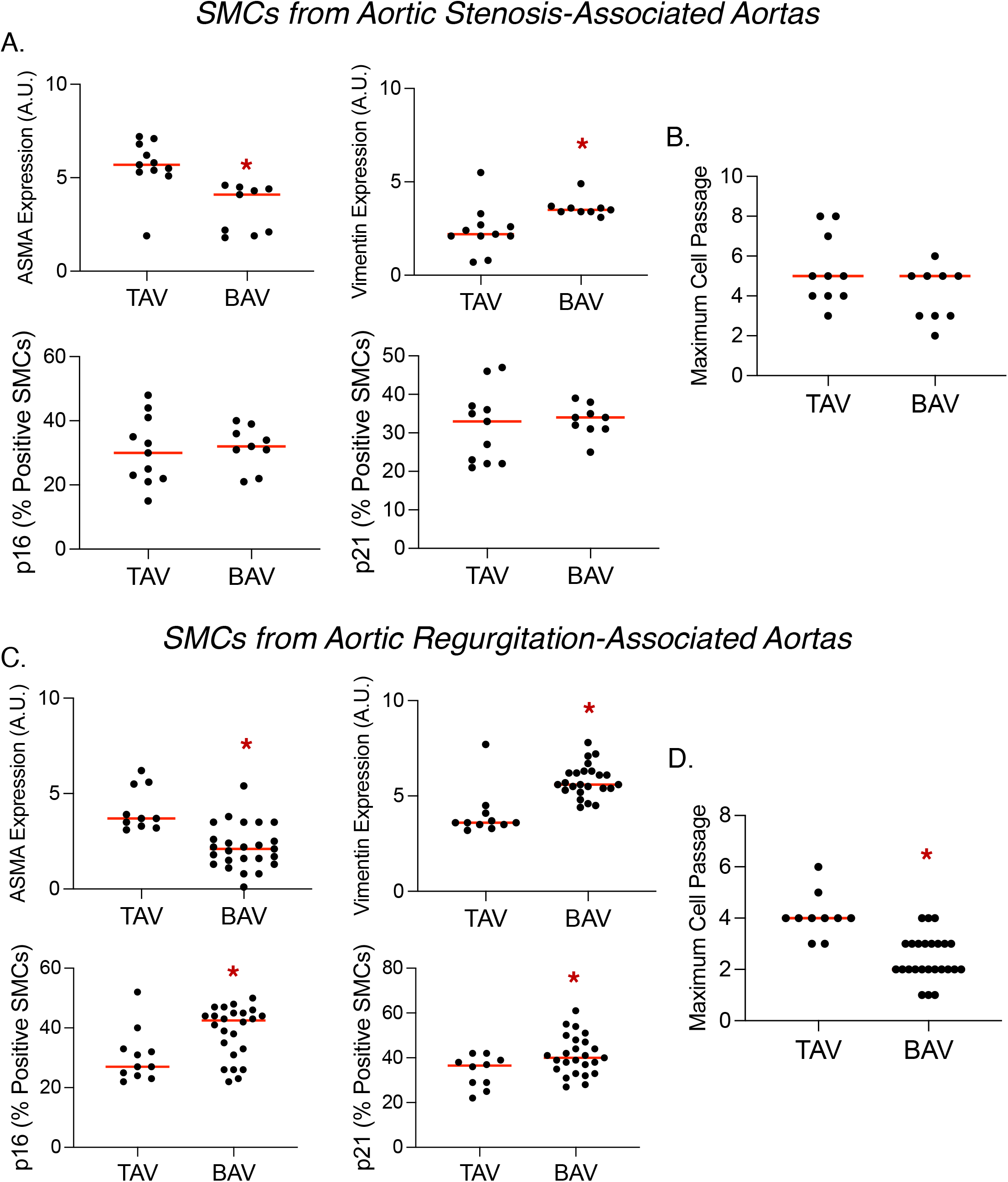
Bicuspid aortic valve (BAV) morphology aggravates phenotypic changes in smooth muscle cells (SMCs) isolated from aortic regurgitation-associated aortas. **A.** Graphs depicting the expression of alpha smooth muscle actin (ASMA), vimentin, p16 or p21 in aortic SMCs from individuals with aortic stenosis and either tricuspid aortic valves (TAV) or BAVs. **B.** Graph depicting the maximum cell passage of cultured SMCs isolated from individuals with aortic stenosis and either TAVs or BAVs. **C.** Graphs depicting the expression of ASMA, vimentin, p16 or p21 in aortic SMCs from individuals with aortic regurgitation and either TAVs or BAVs. **D.** Graph depicting the maximum cell passage of cultured SMCs isolated from individuals with aortic regurgitation and either TAVs or BAVs. Horizontal bars represent median values. *=statistical significance.

Among patients with predominant AR, the expression of ASMA was decreased in SMCs from BAV-AR aortas compared to TAV-AR aortas (*P*<0.0001; **Figure 6c**). Vimentin expression was increased in BAV-AR SMCs compared to TAV-AR SMCs (*P*<0.0001; **Figure 6c**). As opposed to AS, valve morphology appears to have influenced AR-associated cellular senescence, as the expression of p16^INK4a^ and p21^Cip1^ were significantly increased in BAV-AR SMCs compared to TAV-AR SMCs (*P*=0.02 and 0.03, respectively; **Figure 6c**). Moreover, the replicative capacity of BAV-AR SMCs was significantly reduced compared to that of TAV-AR SMCs (*P*<0.0001; **Figure 6d**). Therefore, despite being younger in age, SMCs from patients with BAV-AR exhibit accelerated senescence compared to TAV-AR SMCs.

## Discussion

It has long been recognized that AR has prognostic implications in diseases with ascending aortic involvement. With Marfan syndrome, for instance, the presence of AR increases the risk of aortic dissection and rupture^4, 21^. With BAV and AR, the rate of aortic dilatation and the risk of aortic events is higher compared to BAV with AS^13^ or normally functioning BAVs^12^. Interestingly, these pathological consequences remain even after aortic valve replacement^22, 23^. These findings suggest that an underlying intrinsic factor involving both aortic degeneration and AR may be present in ascending aortic aneurysm disease. In the setting of the aneurysmal aorta, however, it is difficult to determine whether AR has an independent prognostic effect, or whether AR and aortic degeneration are separate entities of a common origin. This is even more complex in the presence of a BAV, in which genetic factors and turbulence are assumed to contribute to aortic degeneration.

We have recently shown AR-specific ascending aortic degeneration, independent of aortic dilatation, aortic valve malformations, or known connective tissue disorders^14^. The most profound degenerative alterations were localized in the medial layer, which included elastin degradation and decreased fibrillin and collagen expression. To this point, the underlying mechanisms have not been investigated. SMCs in the aortic media regulate the turnover of elastin and ECM proteins^24^. Thus, these degenerative findings raise the question of whether SMC-mediated changes drive degeneration in the ascending aortic wall in individuals with AR. SMCs are known to exist in different phenotypes, which relate to different functional properties^25^. We therefore analyzed SMC phenotypes in the non-dilated aorta, as switching from the contractile to synthetic or senescent states primes SMCs for ECM remodeling^6, 26, 27^

Phenotypic switching of SMCs to a synthetic state has been observed in isolated SMCs^28^ and tissue samples from aneurysmal ascending aortas^29–31^. Synthetic SMCs are capable of secreting ECM-degrading enzymes^26, 27^ and may, thus, play a role in medial degeneration in the aneurysmal aorta. We recently found a similar phenotypic switch of SMCs in the normal ascending aorta, which was related to increasing patient age, but independent of aortic dilatation^32^. Here, phenotypic modulation of SMCs was observed in association with AR, and again independent of dilatation. Such changes were minimal with AS and absent in the case of no discernable aortic valve disease. In line with our previous report^32^, phenotypic modulation was more pronounced with increasing age. BAV morphology further aggravated phenotypic modulation, both with AS, and even more so with AR. Interestingly, the BAV patients were significantly younger than the TAV patients in the AR group, suggesting that BAV morphology may be more pathogenic than age. We observed similar phenotypic changes in isolated SMCs, demonstrating that aortic valve disease-related SMC alterations are retained in culture.

SMC differentiation can be influenced by signals from the ECM and intimal endothelial cells^33^. By isolating SMCs from the aortic wall, we were able to clarify that our results are SMC-specific, and independent of recurrent influences of surrounding aortic wall components. Taken together, these findings suggest that AR, patient age and BAV morphology may independently or codependently influence SMC phenotypes in the ascending aorta, prior to dilatation.

More recently it was found that SMCs may enter an irreversible state of cell cycle arrest known as senescence in response to certain injury-related triggers, like unrepaired DNA damage and overwhelming oxidative stress^19^. Although this particular phenotypic switch is thought to be a protective mechanism, senescent SMCs have the ability to compromise surrounding tissue through their senescence-associated secretory phenotype and can reside in the aorta for an extended period of time^6, 20^. Senescent SMCs have been identified in aneurysmal aortas from mice^34^ and humans^6^. In this study, we show that senescence is also associated with AR in the non-dilated ascending aortic wall, suggesting that SMC senescence does not rely on aortic dilatation. We also found that cultured SMCs isolated from AR-associated aortas entered senescence earlier than those from AS-associated aortas, confirming the specificity of a pathogenic AR-effect. It is important to note that SMC senescence was increased in both AS and AR aortas in the presence of a BAV. This is in line with a previous study which showed that SMCs isolated from BAV-associated aneurysms had significantly shorter telomeres than that of TAV aneurysms^35^. Whether factors underlying AR and BAV independently impact SMC senescence, or whether there is interplay between the two pathogenic factors remains to be elucidated.

The findings herein confirm for the first time that AR has a negative impact on the ascending aortic wall, independent of aortic diameter and aortic valve morphology. We showed that AR itself negatively impacts SMC phenotype, such that contractile SMCs are replaced by either synthetic or senescent SMCs that may be primed for ECM degradation. Based on our findings, the mechanisms underlying AR-associated phenotypic modulation should be further evaluated; it might be considered as a therapeutic target in the future to prevent or delay ascending aortic aneurysm formation in patients with AR. Furthermore, we found that SMCs isolated from surgical samples retain their AR-related phenotypic properties in culture. Therefore, SMC isolation from surgical aortic samples provides a medium for studying phenotype-targeted therapeutics.

